# HDAC inhibition ameliorates cone survival in retinitis pigmentosa mice

**DOI:** 10.1101/2019.12.13.874339

**Authors:** Marijana Samardzija, Andrea Corna, Raquel Gomez-Sintes, Mohamed Ali Jarboui, Angela Armento, Jerome E. Roger, Eleni Petridou, Wadood Haq, Francois Paquet-Durand, Eberhart Zrenner, Günther Zeck, Christian Grimm, Patricia Boya, Marius Ueffing, Dragana Trifunović

**Author notes:** Marijana Samardzija, Andrea Corna and Raquel Gomez-Sintes should be considered as shared first authors. EP’s present address is: Institute of Neuronal Cell Biology, Technische Universität München, München, Germany. Corresponding author: Dragana Trifunović, Institute for Ophthalmic Research, Eberhard Karls Universität Tübingen, Elfriede-Aulhorn-Strasse 7, 72076 Tübingen, Germany.

## Abstract

Cone photoreceptor cell death in inherited retinal diseases, such as Retinitis Pigmentosa (RP), leads to the loss of high acuity and color vision and ultimately to blindness. In RP, a vast number of mutations perturb the structure and function of rod photoreceptors while cones remain initially unaffected. Cone death follows rod death secondarily due to increased oxidative stress, inflammation, and loss of structural and nutritional support provided by rods. Here, we show that secondary cone cell death in animal models for RP was associated with an increased activity of histone deacetylates (HDACs). A single intravitreal injection of an HDAC inhibitor at a late stage of the disease, when majority of rods have already degenerated, was sufficient to delay cone death and support long-term cone survival. Moreover, the surviving cones remained light sensitive and initiated light-driven ganglion cell responses. RNA-seq analysis of protected cones demonstrated that HDAC inhibition led to multi-level protection *via* regulation of different pro-survival pathways, including MAPK, PI3K-Akt, and autophagy. This study suggests a unique possibility for a targeted pharmacological protection of both primary degenerating rods and secondary dying cones by HDAC inhibition and creates hope to maintain vision in RP patients independent of the disease stage.

## Introduction

Blindness is a devastating condition, particularly so if it begins in early adulthood as it happens in certain hereditary diseases of the retina characterized by photoreceptor degeneration, such as in Retinitis Pigmentosa (RP). In RP, the leading cause of inherited blindness, mutations in more than 90 genes affect survival and function of rod photoreceptors or retinal pigment epithelium cells (RPE) (1, 2) (http://www.sph.uth.tmc.edu/Retnet/home.htm). One of the particularities of the disease is that despite mutation-unaffected, cone photoreceptors die secondarily once most rods are lost (3, 4). In humans, loss of rods initially has only minor consequences for vision and the majority of patients are unaware of their condition until they start experiencing a prominent reduction in the central visual field, acuity or color discrimination, due to the loss of cones. Hence, in a clinical setting, it is highly pertinent to develop therapies for the treatment of advanced stages of RP, when majority rods have already degenerated and cone degeneration has set in (5, 6).

Despite the importance of cones for human vision, studies on therapeutic options to prevent their loss at advanced stages of RP are disproportionally low (3, 7–11). The reason for this could lie in intercellular relationship between rod and cone photoreceptors in human and mouse retina, where cones account for less than 5% of all photoreceptors (12). Moreover, current knowledge suggests that the massive loss of rods in the late RP creates a “point of no return” after which cone cell death is unstoppable (6, 13). When all rods have degenerated, cones are suffering from the loss of structural and nutritional support from rods (3, 14, 15), exposure to oxidative stress (16) and inflammation (7, 11). Although, alleviating each of these processes individually has the potential to preserve cones to some extent (3, 7, 9, 10, 14), an ideal therapeutic option should provide multi-level protection of cones in the rod depleted retina. One way to achieve this could be by an epigenetically driven simultaneous regulation of a number of genes that are involved in diverse pro-survival responses.

Histone deacetylases (HDACs) are regulators of the chromatin structure and changes in their activity affect transcription of a number of genes (17, 18). Tightly packed chromatin, following the HDAC-governed removal of acetyl groups from histones, is generally associated with transcriptional silencing, albeit this largely depends on the type and “health” status of cells (19–21). Aberrant HDAC activity is causatively linked to various diseases ranging from cancer, through muscular dystrophies, to neurodegenerative diseases (22, 23). We and others have previously shown that regulation of epigenetic patterns, via HDAC inhibition can protect primary degenerating photoreceptors in inherited retinal dystrophies caused by mutations in different genes (24–28). Consequently, more than 90 clinical trials involving HDAC regulators stress HDAC inhibition as promising therapeutic approach for various diseases including retinal dystrophies (21, 29).

Here, we investigated the involvement of HDACs in secondary cone degeneration in mouse models of RP. We found an increased HDAC activity present in both mutation-affected rods and in secondary dying cones. To assess the therapeutic potential of HDAC inhibition for the prevention of cone degeneration in RP, an HDAC inhibitor, Trichostatin A (TSA) was applied at an advanced stage of retinal degeneration, where essentially only a limited number of structurally collapsed cones were left. A single intravitreal injection afforded a long-term preservation of cone photoreceptors. Prolonged cone survival was accompanied by an increase in functional responses. Transcriptional changes associated with cone survival comprised regulation of distinct pro-survival mechanisms, including autophagy, MAPK and PI3K/Akt regulation. Therapies based on HDAC inhibition thus can offer a unique possibility to attenuate loss of photoreceptors independent of the stage of degeneration.

## Results

### Secondary cone degeneration in RP is associated with an increased HDAC activity

To assess the involvement of HDACs in the secondary cone degeneration, we used an HDAC *in situ* activity assay (26) at advanced stages of photoreceptor degeneration in two mouse models for RP, the *rd1* and *rd10* mice. Different mutations in the rod specific phosphodiesterase 6b (*Pde6b*) gene lead to fast rod photoreceptor degeneration, with the onset at post-natal day (PN) 9 in *rd1* and PN14 in *rd10* mice (4, 25, 30). In both animal models cone degeneration begins once most rods have degenerated, around PN20 in the *rd1* and PN40 in the *rd10* mouse (3, 7). At PN30, the outer nuclear layer (ONL) in *rd1* mice is reduced to only one row of photoreceptors, almost exclusively cones (Figure 1A). To localize HDAC activity within the ONL, we combined the *in situ* HDAC activity assay with cone- and rod-specific immunostaining. In line with our previous results HDAC activity was detected in degenerating rods (Figure 1A) (25, 26). Interestingly, HDAC activity also colocalized with the secondary degenerating cones (Figure 1B). The presence of HDAC positive cells within the ONL was detected through the entire period of cone cell death in *rd1* mice (Figure 1B), while in no positive signal for HDAC activity was detected in nuclei of wildtype mice at PN14 (24) and PN30 (data not shown). These findings suggested that the cell death of both mutation-affected rods and secondary dying cones, involves HDAC overactivation. Along the same line, late stages of *rd10* photoreceptor loss also included an increased HDAC activity (Figure 1C, D). The time line of photoreceptor cell death in the *rd10* mouse showed an early peak corresponding to the massive rod loss up to PN26 (30) (Figure 1D). Following, a continuous photoreceptor cell death was detected, likely reflecting the simultaneous rod and cone loss. Similarly to the *rd1* mouse, an increase in HDAC activity paralleled photoreceptor degeneration. The *in situ* assay in retinas from two months old *rd10* mice revealed HDAC activity within a cone arrestin-labeled cone (Figure 1C), linking secondary cone degeneration to aberrant HDAC activity also in *rd10* mice.

**Figure 1.**
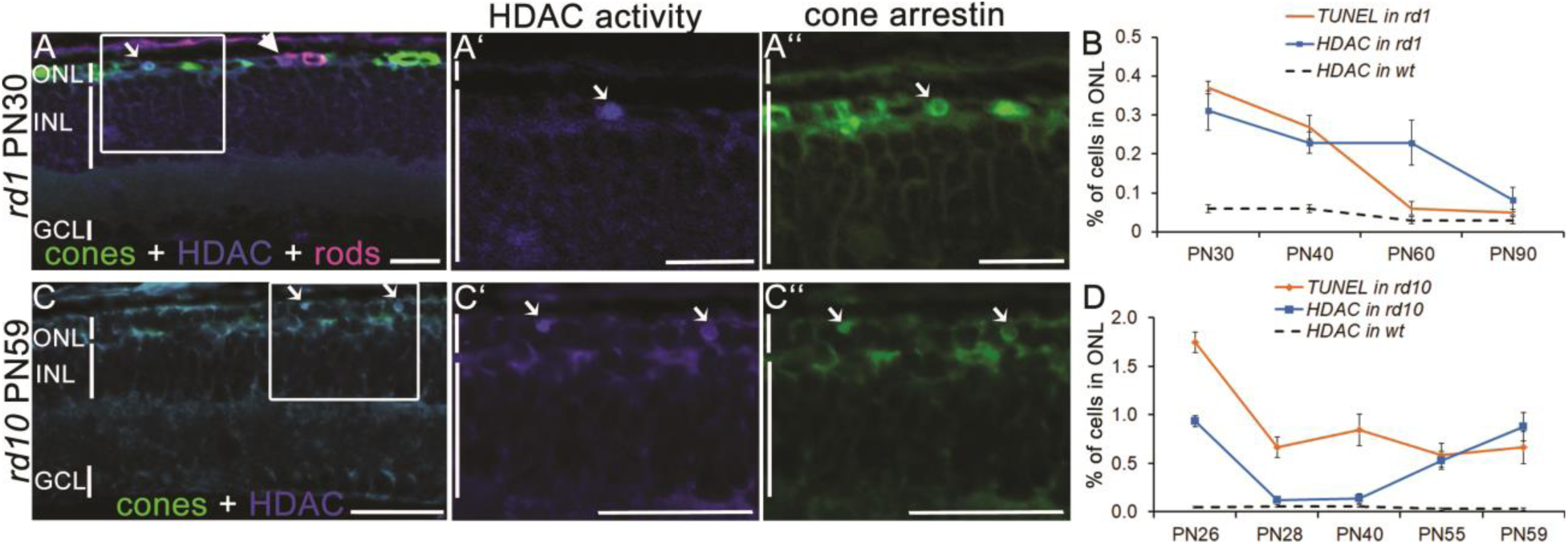
HDAC activity in rd1 retinas at advanced stages of photoreceptor degeneration. (A-A’’) HDAC in situ activity assay in rd1 retinal cross-sections at PN30. HDAC activity (blue) was present in both rod (magenta, arrowhead, rhodopsin immunostaining) and cone photoreceptors (green, arrow, cone arrestin immunostaining). (A’) Magnification of the marked region indicating HDAC+ cell colocalizing with cone arrestin (A’’). (B) Percentage of TUNEL+ and HDAC+ cells in the ONL of rd1 and wt mice over time. (C-C’’) Representative image of a colocalization of HDAC activity and cone arrestin in rd10 mice at PN59, with only cones remaining in the ONL. (D) Percentage of TUNEL+ and HDAC+ cells in the ONL of rd10 and wt mice over time. Data are shown as mean ± SEM (n = 3-5 animals per age). Scale bars: 50 µm. ONL, outer nuclear layer; INL, inner nuclear layer; GCL, ganglion cell layer; PN, post-natal day.

### HDAC inhibition prolongs cone survival in rd1 mice

To determine if HDAC inhibition has a potential to prevent secondary cone degeneration, we injected a well-established class I and II HDAC inhibitor, TSA in the *rd1^TN-XL^* mouse. *rd1^TN-XL^* transgenic mice are homozygous for the mutant *Pde6b* allele and express the fluorescent TN-XL biosensor exclusively in cones, enabling their immediate visualization (31). We have previously shown that a single intravitreal TSA injection prevents mutation-induced cone loss in the *cpfl1* mouse up to 10 days post-injection (24). Additionally, HDAC inhibition delays also primary rod degeneration in both *rd1* and *rd10* retinal explants (25, 26). To minimize the indirect positive effects of increased rod survival afforded by HDAC inhibition on the *rd1* cone survival, we selected a late time-point for the start of the treatment, PN19. Already from PN19, the *rd1^TN-XL^* ONL is reduced only to one row of photoreceptors, almost exclusively cones (Figure 2A). After a single intravitreal injection of TSA at PN19, the effects of HDAC inhibition on cone survival were assessed up to 3 months of age at different time points: PN26, PN30, PN37, PN45, PN60 and PN90. Whole-mount preparations of sham-injected eyes showed center to periphery gradient of cone loss at PN30 (Figure 2B). In contrast, an increased cone-specific fluorescent signal was detected in treated contralateral eyes (Figure 2C). At PN60 loss of cones had proceeded in control eyes, with cones remaining only at the far periphery (Figure 2D). TSA-treated eyes showed higher immunofluorescent signal, suggesting enhanced cone preservation even 41 days following the treatment (Figure 2E). An increase in cone survival was even more evident on retinal cross-sections (Figure 2B’-E’’, Supplementary Figure 1), where individual cone cell bodies and remaining cone segments could be clearly identified. A higher number of cones in TSA-treated animals was also evident on retinal light micrographs (Figure 2F-I’). While untreated retinas showed reduced density of nuclei, many of which were pyknotic, the treated retina displayed healthier cone morphology with classical heterochromatin distribution (32) (Figure 1F-I’). The number of cones in the treated retina after one week remained at the same level as at the start of the treatment (12.03 ± 1.45 SEM cones/100 µm of ONL length at PN19 in untreated retinas *vs*. 12.39 ± 1.43 cones in the treated retina at PN26) (Figure 2J). In the control retinas cone degeneration continued with ∼ 15% less cones than at the start of the treatment (10.12 ± 0.7 SEM in control retinas at PN26). A plot of cone numbers up to PN90 showed that cone loss in control and treated retinas followed an exponential decay function with a clear separation of the two curves (Figure 2J). Additionally, fitted trend lines showed broader separation of the two curves from PN60 and predicted an X-axis intersection of the treated curve with 16 days delay to the control (PN79 for treated *vs.* PN95 control retinas, Figure 2J). These extrapolations suggest that the TSA treatment not only delays but might slow down secondary cone degeneration as well.

**Figure 2.**
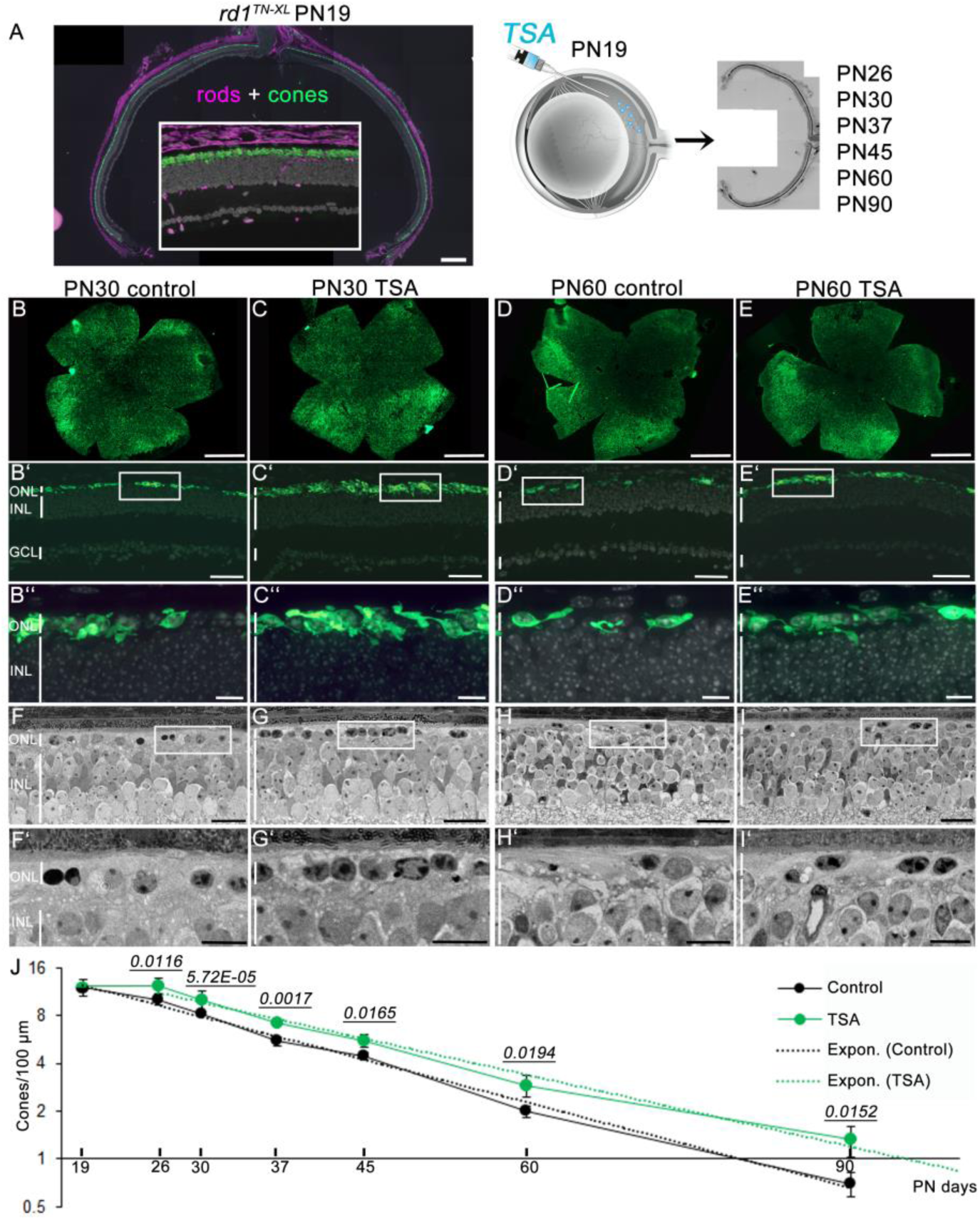
HDAC inhibition promotes long-term cone survival in rd1^TN-XL^ mice. (A) Retinal cross-section of an rd1^TN-XL^ mouse at PN19, showing TN-XL-labeled cones (green), rods (magenta, anti-rhodopsin antibody) and nuclei (grey, DAPI). Note that mouse secondary antibody used to detect anti-rhodopsin antibody showed the non-specific signal in layers other than ONL. Schematic representation of a single intravitreal injection of TSA at PN19 followed by quantification of cone survival up to PN90. (B-D) Flat mounted retinas from control and TSA-treated eyes at indicated PN days. (B’-D’) Representative images of retinal cross-sections from control and TSA-treated retinas at PN30 and PN60 used for quantification of TN-XL labeled cones. (B’’-D’’) Magnifications of the marked regions shown in B’-E’. (F’-I’) Retinal morphology of control and treated retinas at PN30 and PN60. Magnification of marked regions is shown in F’’-I’’. (J) Quantification of cone survival in control and TSA-treated retinas. Y-axis is in log2 scale. Data are shown as means ± SEM (n = 5-10 animals per age). Numerical p values by Mann-Whitney nonparametric test. Fitting of the exponential curves are shown in dotted lines. Scale bars: (A, B’-E’) 50 µm; (B-E) 500 µm; (B’’-E’’, F-I) 20 µm; (F’-I’) 10 µm. ONL, outer nuclear layer; INL, inner nuclear layer; GCL, ganglion cell layer; PN, postnatal days.

To analyze the effects of continuous TSA treatment on cone survival, we treated *rd1^TN-XL^* retinal explants from PN19 until PN26 *ex vivo*. Organotypic retinal cultures, consisting of retina and RPE layer, enable maintaining mature neurons *in situ,* as well as complex neuronal connections, while providing the possibility for constant exposure to a drug via a culture medium (Figure 3A). Similarly to *rd1^TN-XL^* cone degeneration *in vivo*, the center to periphery gradient of cone loss was more prominent in the control explant cultures (Figure 3B). Quantification of cone numbers in retinal cross-sections showed ∼30% increase in the cone survival in TSA-treated retinal explants (Figure 3B).

**Figure 3.**
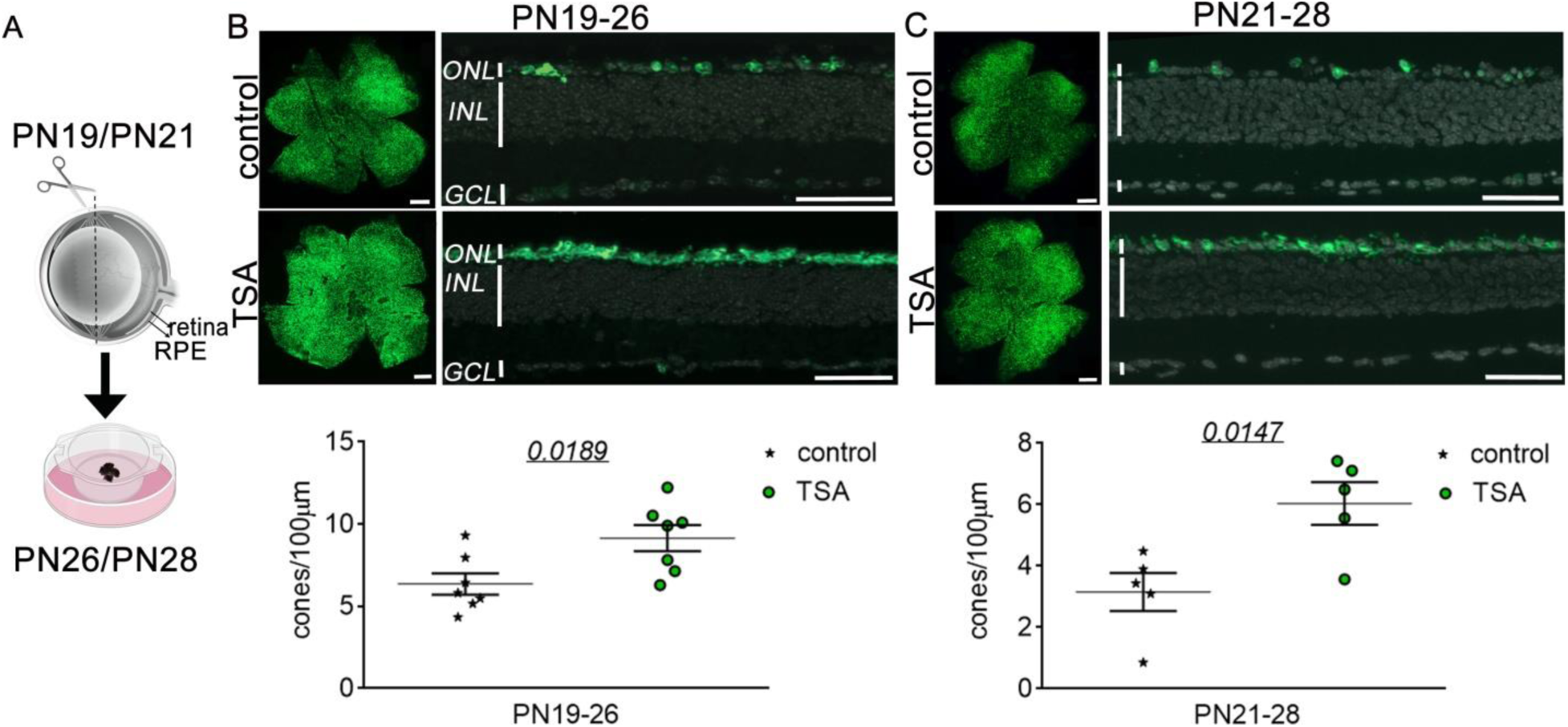
HDAC inhibition protects rd1^TN-XL^ cones ex vivo. (A) Schematic representation of the ex vivo retinal explants preparation. Retinas from rd1^TN-XL^ mice were collected at PN19 or PN21 and treated for 7 days with control or TSA-medium. (B-C) Representative flat mount preparations and cross-sections of explanted retinas that were used for the quantification of cone survival, shown in the dot plot below. Data are shown as mean ± SEM (n = 5-7 animals per age group). Numerical p-values by unpaired, two-tailed t test. Scale bars in whole mounts 500 µm; retinal cross-sections 50 µm. RPE, retinal pigment epithelium; ONL, outer nuclear layer; INL, inner nuclear layer; GCL, ganglion cell layer; PN, postnatal day.

Next, we asked if the HDAC inhibition has a potential to protect secondary dying cones even at a later stage of degeneration. For this, we started the treatment at PN21 and assessed the *rd1^TN-XL^* cone survival after one week in culture (Figure 3C). Also in this case, the TSA treatment significantly improved cone survival, with 52% more cones in the treated explants (6.03 ± 0.69 cones/100 µm of length in the TSA-treated explants *vs.* 3.16 ± 0.62 SEM in the controls, n=5, *p=0.0147*). We also tested the neuroprotective properties of Panobinostat, a clinically approved pan-HDAC inhibitor within the same group of inhibitors as TSA (33, 34). Similarly to TSA, Panobinostat treatment starting at PN19, increased *rd1^TN-XL^* cone survival *ex vivo* up to 30% after 7 days (Supplementary Figure S2).

### HDAC inhibition improves cone-mediated light-responses

Next, we asked whether the remaining cones were light-sensitive and able to drive functional responses. To address the functionality of the TSA-protected cones, we used a micro-electrode array (MEA) (35, 36) to record the light-mediated spiking activity of retinal ganglion cells (RGCs) of *rd1^TN-XL^* retinal explants. The experimental set up included a light source at the bottom of the MEA chamber, mimicking the physiological situation where light stimulation comes from the ganglion cell side, while the RPE layer on the top of explants provided a physiological environment for light absorption (Figure 4A).

**Figure 4.**
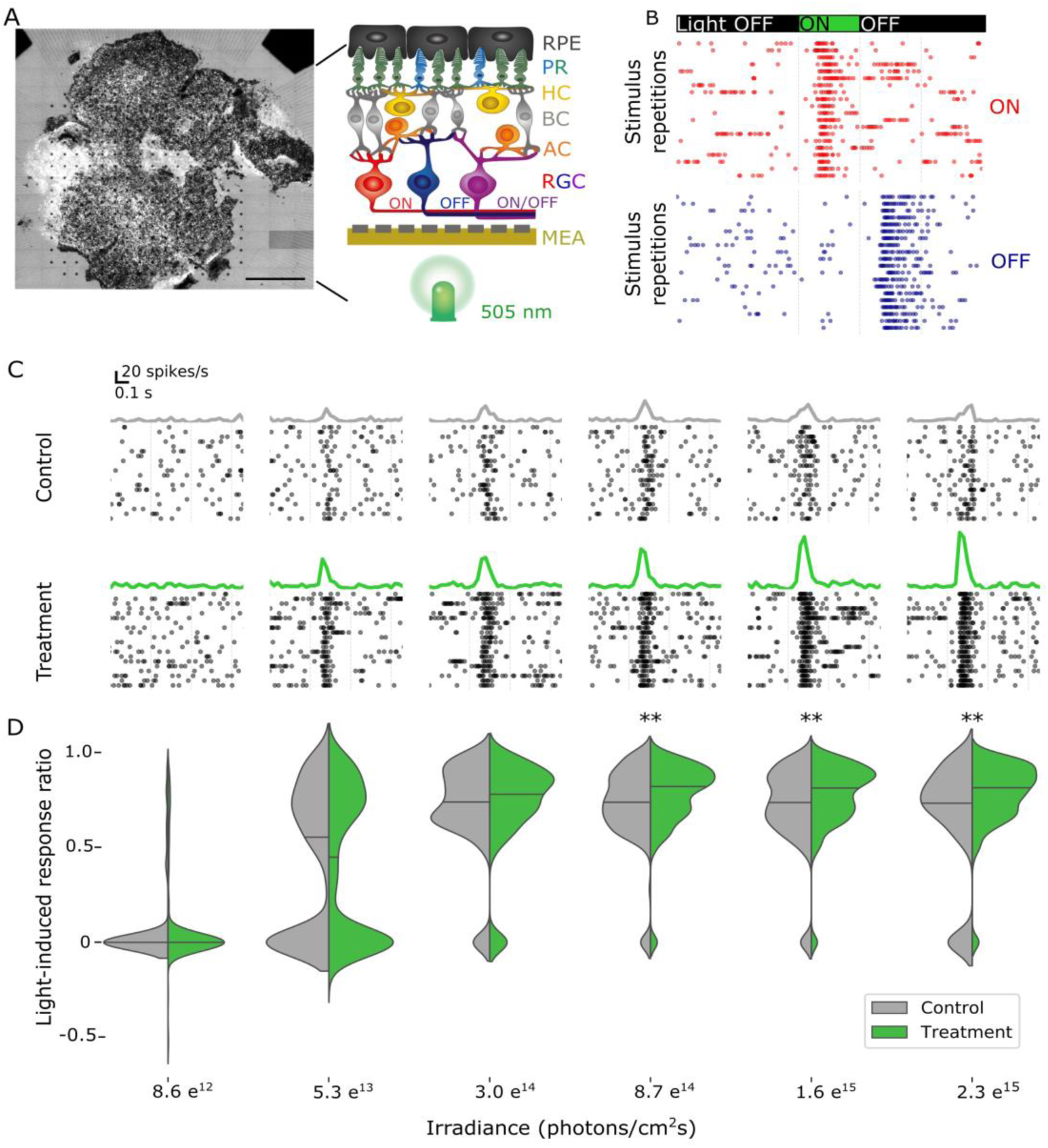
HDAC inhibition improves cone function in rd1^TN-XL^ retinal explants. (A) A micro-electrode array (MEA) with 256 electrodes was used to record retinal ganglion cell (RGC) responses in control and TSA-treated PN19-26 rd1^TN-XL^ retinal explants. Schematic drawing of the MEA setup used in the study. (B) Representative recordings of ON and OFF responses from TSA-treated retinal explants during stimulation with flickering light (350 milliseconds flashes, 505 nm) shown as raster plots. Each dot indicates one spike. Green bar indicates start of light stimulation. (C) Exemplary spike responses obtained in control (gray) and treated (green) explants at six different light intensities, shown as raster plots (bottom) and averaged firing rate histograms (top). (D) Quantification and discrimination of the response ratio in control (grey) and treated explants (green). Significant differences based on Wilcoxon rank-sum test are detected for three high light intensities (n = 161 channels for control, n= 274 for treated retinas, ** p < 0.01). RPE, retinal pigment epithelium; PR, photoreceptors; HC, horizontal cells; BC, bipolar cells; AC, amacrine cells; RGC, retinal ganglion cells.

A LED emitting green light (505 nm) with increasing light intensities was used to stimulate cones in both control and TSA-treated PN19-26 retinal explants. Light-ON and Light-OFF responses were detected in both control and treated retinas suggesting preservation of physiological network functionality (Figure 4B). To estimate the degree of activity change upon light stimulation, we calculated a light-induced response ratio for ON responsive MEA electrodes. The light-induced response ratio (see Method section) quantifies how much the firing rate (Figure 4C) increases during light stimulation and can therefore be considered as a simple link between number of rescued cones and functional readout. We evaluated a total of 161 active ON channels in eight control retinas and 274 channels in nine treated retinas. The distribution of the light-induced response ratio, presented as a violin plot, demonstrates that increasing light intensity lead to a higher firing rate (Figure 4D). Changes of the spontaneous activity under same light intensities were not detected. For each condition (control *vs.* treatment) the light-induced response ratio of the two distributions were compared for statistical difference. While for low intensities the median values in control and treated retinas were not significantly different, the light-induced response ratio increased more strongly in the treated samples and was significantly different above a light intensity of 8.7 e^14^ photons/cm^2^ sec (p < 0.01). The median light-induced response ratio in treated retinas reached 0.82 while in control condition it approached 0.74 (Figure 4D).

### HDAC inhibition induces global changes in gene-transcription patterns in the surviving cones

To gain an insight into molecular signatures governing the TSA-induced cone survival, we performed whole transcriptome analysis of flow-sorted *rd1^TN-XL^* cones of retinal explants after seven days in culture (PN19-26). TN-XL+ cells represented 95% of FASC-sorted cells, suggesting a highly purified cone population (Supplementary Figure S3). RNA-seq analysis of differentially expressed genes (DEG) in TSA-treated *vs.* untreated cones revealed 1845 genes with significantly different expression (enrichment of 1163 and reduction in 682 genes), as indicated in the volcano plot (Figure 5A). The complete list of differentially expressed genes is provided in Supplementary table 2. Regulation of signaling pathways can be influenced by coordinated subtle changes in proteins abundance and modifications, rather than the level of expression of individual genes (37, 38). As RNAseq data provide information about gene expression levels that may not necessarily influence protein abundance (39), we focused our analysis on changes in pathways modulation. Therefore, for the pathway analysis we used protein-coding genes with significant differential expression following the TSA treatment (p≤ 0.05), regardless of their fold change. This enabled identification of signature trends for activation or inhibition of downstream pathways. Top molecular pathways differentially regulated following the TSA treatment, included transcriptional changes of around 40 genes belonging to PI3K-Akt and/or MAPK signaling pathways and more than 20 genes regulating cellular senescence, endocytosis, actin cytoskeleton, as well as cAMP and calcium signaling (Figure 5A, PaintOmics link: http://www.paintomics.org/?jobID=4CIbSr1eA4). As the RNA-seq analysis suggested changes in PI3K-Akt and MAPK signaling, both of them extensively linked to neuroprotection (40, 41), we retrieved genes belonging to each of these pathways from the KEGG database. The heatmap representation of their expression demonstrated the enrichment of about half of the genes (23/42) belonging to the PI3K-Akt (mmu04151) and to the MAPK cascade (20/36, mmu04010) in the TSA-treated cones (Figure 5B). Among highly abundant transcripts were the brain-derived neurotrophic factor (*Bdnf*) and the insulin-like growth factor 1 (*Igf1*), growth factors regulating MAPK and PI3K-Akt pathways. Significantly increased *Bdnf* expression in the TSA-treated cones sorted from independent samples was confirmed by quantitative PCR (qRT-PCR) (Figure 5C).

**Figure 5.**
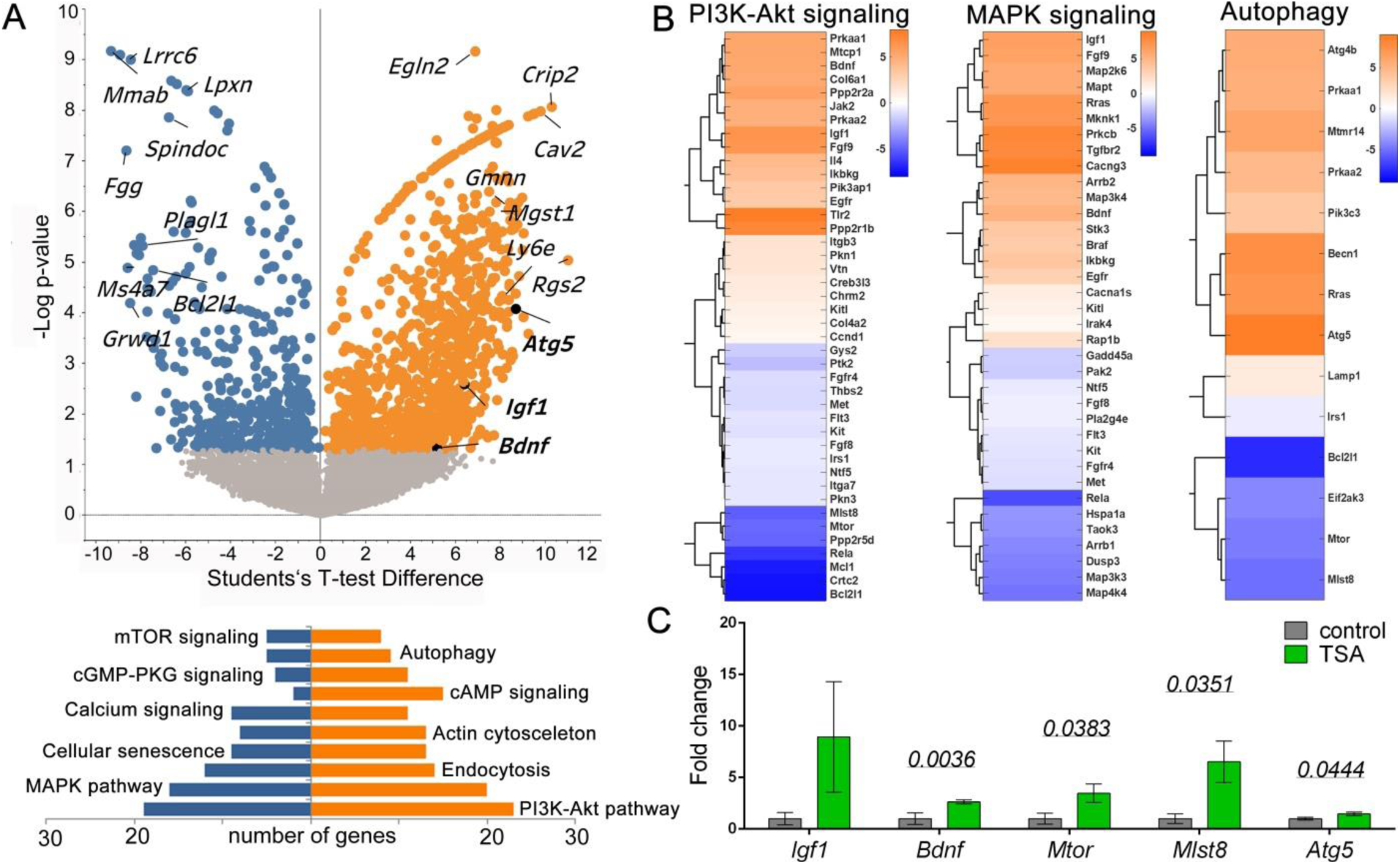
Whole genome transcriptomic analysis of TSA-induced survival of rd1^T-NXL^ cones. (A) Volcano plot representation of differentially expressed genes as detected by RNA-seq in flow-sorted cones from treated vs. untreated PN19-26 rd1^TN-XL^ (n = 3 animals). Orange and blue dots: significantly enriched and perturbed genes, respectively, in TSA treated cones (FDR-based, Student’s T-test of mean difference). PaintOmics analysis was used to identify differentially regulated pathways in TSA-protected cones. (B) Heat maps with hierarchical clustering of differentially expressed genes within PI3k-Akt, MAPK and autophagy pathways. Genes showing Student’s t-test difference between TSA and control, with p≤ 0.05 were selected. (C) qRT-PCR validation of differential expression of PI3K-Akt, MAPK and autophagy regulators in FACS-sorted cones. Fold changes are relative to controls. Data are shown as mean ± SEM (n = 4 animals). Numerical p-values by Mann-Whitney nonparametric test.

Moreover, the pathway analysis showed significant upregulation of most of the genes (9/14) related to autophagy (mmu04140), an additional pro-survival mechanism in neurodegenerative diseases (Figure 5B) (42, 43). Upregulation of the autophagy-related-5 gene (*Atg5*) in surviving cones was confirmed by qRT-PCR (Figure 5C). While, RNAseq data suggested downregulation of *Mtor* and *Mlst8*, regulators of both MAPK and autophagy pathways, qRT-PCR on independent samples showed significant upregulation in the treated cones. To evaluate if increased *Atg5* transcription was resulting in increased autophagy, we assessed formation of autophagolysosomes in surviving cones. To this end, we quantified the colocalized puncta between an autophagosome marker, the microtubule-associated proteins 1A/1B light chain 3B (LC3B) and lysosome-associated membrane protein 1 (LAMP1) in control and treated *rd1^TN-XL^* cones. LC3-LAMP1 staining of retinal cross-section from PN19-26 *in vivo* TSA-treated mice showed overall increase in both LC3- and LAMP1-positive vesicles, as well in colocalizing puncta within the cones in comparison to sham-treated retinas (Figure 6). Similarly, increased LC3-LAMP1 colocalization in cones was also detected in *ex vivo* treated *rd1^TN-XL^* explants (Supplementary Figure S4).

**Figure 6.**
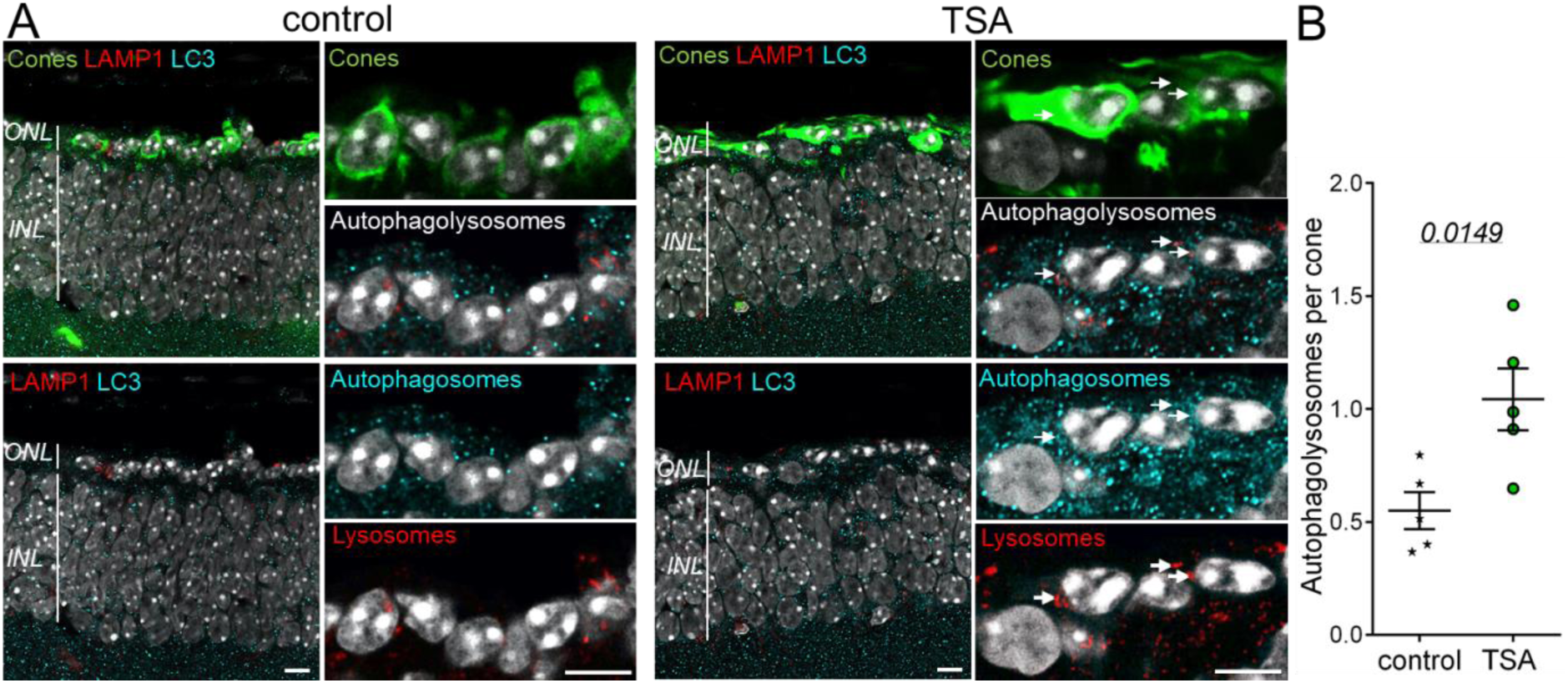
Autophagy after HDAC inhibition in vivo. (A) Representative images from a single confocal plane on one retinal cross-section from a rd1^TN-XL^ mouse at PN26, sham- or TSA-injected at PN19, with TN-XL cones (green), autophagosomes (cyan, LC3B antibody) and lysosomes (red, LAMP1 antibody). Colocalization between autophagosomes and lysosomes is marked with arrows. (B) Quantification of the number of colocalized LC3-LAMP1 (autophagolysosomes) puncta per cone in the ONL. Data are shown as mean ± SEM (4 positions within retina, n = 3 animals). Numerical p-values by Mann-Whitney nonparametric test. Scale bars 10 μm. ONL, outer nuclear layer; INL, inner nuclear layer.

Finally, to identify similar molecular signatures to the expression pattern obtained in this study, we submitted our data to the Library of Integrated Network-Based Cellular Signatures project, (LINCS) (Supplementary Figure S5) (44). The analysis matched our data to gene expression profiles of TSA and Vorinostat treatments on human cell lines deposited in the LINCS platform. Of note, Vorinostat is a clinically approved pan-HDAC inhibitor from the same class as TSA. This suggests a possibility for the repurposing of “off the shelf” HDAC-inhibiting drugs for the treatment of RP.

## Discussion

Cone photoreceptors are indispensable for human vision in daylight and their loss in inherited or acquired retinal diseases have devastating effects on daily tasks and life quality (27, 45, 46). In RP, mutations in any of 90 identified genes will not only lead to loss of rods, but will ultimately result in cell death of cones despite them being not directly affected by mutations (47, 48). Dependence of cones on rods’ survival is considered to be crucial, as loss of 95% of all photoreceptors will expose the remaining cones to oxidative stress, inflammation, nutritional deficit, as well as loss of the structural and trophic support normally provided by rods (3, 7, 14-16). However, despite such adverse environmental conditions and loss of their functionality, cones can persist for a long time in RP patients. Dormant cones retain cell bodies long after the loss of outer and inner segments, as well as a potential for structural and functional revival under certain conditions, such as optogenetic reactivation or upon transplantation of rods (49, 50).

Here, we show that cones also have an innate ability for survival in the total absence of rods, which can be induced by HDAC-driven transcriptional changes. At the same time HDAC inhibition may affect other cells in the retina, which could support cone survival in the absence of rods (51, 52). A single intravitreal injection of HDAC inhibitor fully protected cones for one week and significantly slowed down cone loss up to 90 days even when all rods had already degenerated. Interestingly, the *rd1^TN-XL^* cone loss followed exponential kinetics as already suggested for other forms of neurodegeneration (53). The observed exponential cone loss in the *rd1^TN-XL^* retina excludes the possibility of cumulative damage caused by the loss of rods, and suggests that secondary dying cones can be rescued at any time, although with the disease progression fewer cells will be amenable for rescue. Therefore, a treatment even at late stages of disease is likely to be beneficial (53). Indeed, our data on HDAC-induced cone protection at different stages of *rd1^TN-XL^* cone loss (PN19 and PN21) confirm this and is furthermore in line with other observations indicating that the window-of-opportunity for the treatment of retinal dystrophies is much broader than it is currently considered (Samardzija et al., in press, (6)).

Preserving retinal function in the absence of rods is the ultimate goal of neuroprotective therapies for late stage RP. Indeed, recent studies demonstrated some improvements in visual function, following therapies specifically targeting cones in mouse models of RP with slower degeneration and at stages where most of rods are still present (7, 9). However, the low number and the impaired morphology of the remaining cones at the stages where rods have already fully degenerated is a major obstacle in assessing visual improvements by standard functional tests like ERG (54). We did not detect changes in ERG-wave amplitudes in *rd1^TN-XL^* mice treated at such late stages of degeneration (data not shown). However, by recording cone-mediated light-induced responses from large populations of individual RGCs we were able to demonstrate intact cone-driven retinal circuitry even in the absence of rods. While a light-induced increase in spiking was observed over 2 log units (55), the treatment did not change the ongoing, spontaneous activity of RGCs. Based on the transient spiking responses to light we can exclude an activation of intrinsically photosensitive RGCs (56) and conclude an overall improvement of light responses upon HDAC inhibition.

For the treatment of retinal diseases intravitreal injections are the preferred route of administration compared to subretinal injections, as larger volumes of drug can be applied and the risk of retinal damage is reduced. However, due to an extensive retinal circulation and the permeability of the brain-retinal-barrier for low molecular weight compounds, intravitreally injected TSA is cleared from the mouse eye in less than two hours (24, 57). Still, a single TSA injection led to a long-lasting cone survival suggesting the possibility of epigenetically induced pro-survival mechanisms that could counteract environmental insults caused by the absence of rods. While HDAC inhibition induces changes in gene expression within minutes (28), the effects of transcriptional regulation can persist for months (58). To get an insight into the protective effects of HDAC inhibition on secondary cone death, RNAseq analysis of continuously treated cones enabled identification of transcriptional changes associated with the HDAC inhibition. At the same time, our data provide a source for further studies into the overarching transcriptional changes facilitating cone survival in the late stages of RP.

In advanced RP cones are exposed to high oxidative stress and inflammation (7, 16, 59). Oxidation induced injury at the cellular level elicits a broad range of responses, from proliferation to cell death, that largely depend on the balance between various cell signaling pathways activated as a response to oxidative stress (60). Our data point to changes in two major signaling pathways involved in regulating cellular response to oxidative stress, the mitogen-activated protein kinases (MAPK) and phosphoinositide 3-kinase activated protein kinase B (PI3K-Akt) pathways. Although MAPK pathways comprise a large number of kinases that can regulate cellular survival in opposing manner, MAPK regulation is linked to increased photoreceptor survival (61–63). German et al., demonstrated protective effects of docosahexaenoic acid (DHA) on photoreceptor survival via MAPK pathway activation (62). In addition, a positive involvement of p38 MAPK was later causally connected to LIF-dependent neuroprotection during photoreceptor degeneration (63). On the other hand, the importance of the PI3K-Akt pathway in photoreceptor survival is well-established, with PI3K-Akt members acting downstream of various neuroprotective agents (64, 65). In addition, the PI3K-Akt pathway connects extracellular signals, such as insulin, with the mTOR pathway shown to play an important role in secondary dying cones (3, 9). RNA-seq data obtained in this study indicate regulation of more than 40 genes within the PI3K-Akt pathway, further confirming the PI3K-Akt pathway as one of the major regulators of cone survival in RP. Similarly, MAPK and PI3K-Akt regulation is also associated with inflammation in various neurodegenerative conditions, including retinal diseases (66, 67). Interestingly, our data suggest that an increase in transcription of neurotrophic factors, including *Igf1* and *Bdnf*, may contribute to cone survival. Although BDNF-induced neuroprotective effects on light- or mutation-induced photoreceptor degeneration was recognized as early as 1992 by LaVail and colleagues (68, 69) our study indicate a direct upregulation of *Bdnf* transcription in the cones protected from secondary degeneration.

Besides increased oxidative stress and inflammation, cone starvation has recently been recognized as one of the main contributors to secondary cone death in RP (3, 9, 14). Autophagy is the main cellular mechanism for self-nourishment and recycling of metabolites to supplement macromolecules and energy under severe starvation (42, 43). The beneficial role of autophagy is demonstrated for the clearance of misfolded proteins in mutation-affected photoreceptors or in dysfunctional RPE in age-related macular degeneration (AMD) (70–73). Yet, in secondary cone degeneration it remained unclear whether increased autophagy is beneficial or not (3, 74, 75). The observed upregulation of *Atg5* transcription and an increased number of autophagolysosomes in TSA-treated cones are in line with the previously reported effects of HDAC inhibition on autophagy induction (76) and highlight autophagy as a protective mechanism in secondary cone degeneration. Interestingly, while RNA-seq analysis of transcriptional regulation following the TSA treatment suggested differential regulation of both autophagy and mTOR pathways, *via* downregulation of *Mtor* and *Mlst8* among other genes, qRT-PCR validation showed significant upregulation of both transcripts. Whether the observed differences in the TSA-induced transcriptional regulation of *Mtor* and *Mlst8* will induce an activation of the mTOR pathway, associated with an increased cone survival in RP (3, 9), requires further studies.

Taken together, our results demonstrate multi-level and long-lasting effects of HDAC inhibition in the prevention of late stages of secondary cone degeneration in a mouse model for RP characterized by fast photoreceptor loss. In combination with our previous data linking HDAC overactivation with the loss of rods in eight different mouse models of RP (25, 30), this study highlights HDACs as a common denominator of both mutation-induced rod cell death and secondary cone degeneration and represents a unique therapeutic possibility for the treatment of RP independent of the stage of degeneration. Finally, detrimental environmental conditions inducing cone loss at late stages of RP, share important similarities with age-related macular degeneration (AMD), the leading cause of blindness in the industrialized world (77). Although RPE cells are considered as the primary target for AMD pathology, loss of cone photoreceptors in macular region in AMD patients characterize final stages of the disease (78–80). As dysfunction of RPE cells in AMD may expose cones to a milieu similar to the one in late RP, HDAC inhibition holds promise also for the treatment of more common forms of retinal degeneration.

## Methods

### Animals

The C3H *rd1/rd1 (rd1),* C57BL/6J x C3H *HR2.1:TN-XL x rd1 (rd1^TN-XL^)* and C57BL/6J *rd10/rd10 (rd10)* mice were housed under standard light conditions, had free access to food and water, and were used irrespective of gender. *rd1^TN-XL^ mice*, express the TN-XL (Ca^2+^ biosensor) selectively in cone photoreceptors under the control of the human red opsin promoter (HR2.1) (31, 81). The presence of TN-XL biosensor does not alter the *rd1* phenotype, while it enables direct visualization of cone photoreceptors by fluorescence microscopy (81). All procedures were performed in accordance with the ARVO statement for the Use of Animals in Ophthalmic and Vision Research and the regulations of the Tuebingen University committee on animal protection, Germany and veterinary authorities of Kanton Zurich, Switzerland.

### Intravitreal injections

Single intravitreal injections were performed at PN19 in *rd1^TN-XL^* mice that were anesthetized subcutaneously with a mixture of ketamine (85 mg/kg) and xylazine (4 mg/kg). One eye was injected with 0.5 µl of a 100 nM TSA (catalog T8552, Sigma-Aldrich) in 0.0001% DMSO, while the contralateral eye was sham-injected with 0.0001% DMSO and served as a control. Assuming the intraocular volume of mouse eye to be 5µl (82), this procedure resulted in a final intraocular concentration of 10 nM TSA.

### Retinal explant cultures

Organotypic retinal cultures from *rd1^TN-XL^* animals including the retinal pigment epithelium (RPE) were prepared under sterile conditions as previously described (24, 25). Briefly, PN19 or PN21 animals were sacrificed, the eyes enucleated and pretreated with 0.12% proteinase K (ICN Biomedicals Inc.) for 15 minutes at 37°C in HBSS (Invitrogen Inc.). Proteinase K activity was blocked by addition of 10% fetal bovine serum, followed by rinsing in HBSS. In the following, the cornea, lens, sclera and choroid were removed, while the RPE remained attached to the retina. The explant was cut into a clover-leaf shape and transferred to a culture membrane insert (Corning Life Sciences,) with the RPE facing the membrane. The membrane inserts were placed into six well culture plates with Neurobasal-A medium (catalog 10888022) supplemented with 2% B27 (catalog 0080085-SA), 1% N2 (catalog 17502048) and L-glutamine (0.8 mM, catalog 25030032) (all from Invitrogen Inc.), and incubated at 37°C in a humidified 5% CO2 incubator. The culture medium was changed every 2 days during the 7 days culturing period. Retinal explants were treated with 10 nM TSA or 1 µM Panobinostat (catalog S1030, Selleckchem) diluted in Neurobasal-A culture medium. For controls, the same amount of DMSO was diluted in culture medium. Culturing was stopped after 7 days by 2 h fixation in 4% PFA, cryoprotected with graded sucrose solutions containing 10, 20, and 30% sucrose and then embedded in tissue freezing medium (Leica Microsystems Nussloch GmbH).

### Histology

For retinal cross-sectioning, the eyes were marked nasally and eye cups (after cornea, iris, lens and vitreous removal) were fixed in 4% paraformaldehyde for 2 hours at room temperature. Following graded sucrose cryoprotection eyes were embedded in optimal cutting temperature compound (Tissue-Tek), cut into 12 μm sections and mounted with Vectashield medium containing 4’,6-diamidino-2-phenylindole (DAPI, Vector). For retinal flat mounts, retinas without RPE were fixed for 30 minutes, cut into a clover leaf shape and mounted with Vectashield with the photoreceptors facing up. To analyze retinal morphology, eyes were fixed in 2.5% glutaraldehyde, cut at the level of the optic nerve, followed by 1% osmium tetroxide treatment post fixation and ethanol dehydration, according to a previously described protocol (32). After embedding in Epon 812, 0.5 μm thick sections were counterstained with toluidine blue. Immunostaining was performed on retinal cryosections by incubating with rabbit primary antibodies against cone arrestin (1:1000; catalog AB15282, Merck Chemicals GmbH), mouse anti-rhodopsin (1:400, catalog MAB 5316, Chemicon), LC3B (1:100 Novus, cat NB-100-220) and LAMP1 (1:100, clone 1D4B, DSHB) at 4°C overnight. Alexa Fluor 488, 568 or 647-conjugated antibodies were used as secondary antibodies. Images were captured using Z-stacks on a Zeiss Axio Imager Z1 ApoTome Microscope using 20x air, 40x oil or 100x oil objectives. For the quantification of the LC3 and LAMP1 puncta in cones, 4 images per retina for *in vivo* treatment and 8 images per explant, were assessed in each confocal plane obtained by the Leica TCS SP5 Confocal Microscope. Colocalizing puncta were counted using the counter plugin of Image J. The number of colocalizing puncta was divided by the number of cones in the whole z-stack.

### Quantification of cone survival

The quantification of cones was performed by manually counting the number of TN-XL labeled cones (using the Zen event counter) on at least two retinal cross-sections cut along the dorsoventral axis at the level of the optic nerve. Retinal cross-sections were used for quantification of cone photoreceptors as the TN-XL biosensor is present throughout the cone photoreceptor, except the outer segment (IS) (31). The presence of the biosensor in the cell body, axon and IS, hampers clear separation of individual cell bodies from the IS and/or axon on flat mount preparations at the late stages of *rd1^TN-XL^* degeneration, where cones align horizontally to the INL due to the lack of structural support from rods (Supplemental Figure 1A.) Retinal cross-sections with labeled nuclei enabled distinction between different parts of the cones and facilitated the counting of their cell bodies. Cones were quantified on multiple images projection (MIP) obtained from 9-15 optical sections taken with 20x magnification at four positions in the retina: ventral and dorsal central retina (corresponding to −10° and 10° of eccentricity from the optic nerve –ON, respectively) and ventral and dorsal peripheral positions at −80° and 80° degrees (Supplemental Figure 1). Spider diagrams show the number of cones per 100 µm of the outer nuclear layer (ONL) length at each position, presented as mean values ± SEM.

### HDAC in situ activity assay

HDAC activity assays were performed on 12 µm thick cryosections of 4% PFA-fixed eyes following immunostaining against cone arrestin/rhodopsin as previously described (4). Briefly, retina sections were exposed to 200 μM Fluor de Lys-SIRT2 deacetylase substrate (Biomol) with 500 μM NAD^+^ (Biomol) in assay buffer (50 mM Tris/HCl, 137 mM NaCl; 2.7 mM KCl; 1 mM MgCl_2_; pH 8.0) for 3 hours at room temperature. Following methanol fixation at −20°C for 20 min, a developer solution (1x Biomol; KI105) containing 2 μM TSA and 2 mM nicotinamide in assay buffer was applied to generate the signal. Due to the presence of a background staining in negative controls, only cells with a prominent nuclear staining were considered as HDAC positive (24).

### MEA recording

Retina explant cultures attached to membrane were transferred from the incubation chamber to a 256-electrode MEA (Multi channel systems MCS GmbH) with the ganglion cell side facing the electrodes. A custom-made grid was placed over the retina to improve the contact between electrodes and the tissue and the stability of the recording. Cultures were perfused throughout the experiment with oxygenated Ames’ medium (A 1420, Sigma) and heated to 36°C. The electrode spacing was 200 µm with the total recordings area of ∼ 3.2 x 3.2 mm^2^. Twenty repetitions of 350 ms long light-flashes of increasing intensity (8.6 e^12^, 5.3 e^13^, 3 e^14^, 8.7 e^14^, 1.6 e^15^, 2.3 e^15^ photons/cm^2^ s) separated by 2 s of dark were presented to both control and the TSA-treated retinal explants mounted on MEA. The recordings were made with a sampling rate of 25 kHz using the MC Rack software (Multi Channel Systems MCS GmbH). The analysis of recordings from 256-MEAs was performed using Python 3.6. Recordings were bandpass - filtered (400 – 5000 Hz, Butterworth 2nd order) and spikes were detected as threshold crossing of 5 time the standard deviation of the filtered signal with a pause time of 1.5 ms.

Light-induced RGCs activation recorded by an electrode was quantified for each light intensity using a two tailed, paired *t-test* comparing the detected spikes during the 20 repetition of light-ON (350 ms of light flash) or light-OFF (350 ms after the light shut-off) versus the spontaneous activity recorded before light onset. Only electrodes with a statistically significant difference (p<0.01 and t-statistic > 2) were considered as light-activated. In addition, only channels light-activated for at least 3 out of 5 light intensities in the photopic light range (5.3 e^13^, 3 e^14^, 8.7 e^14^, 1.6 e^15^, 2.3 e^15^ photons/cm^2^) were included in the analysis. These criteria were used to eliminate potential non-stable recordings. To quantify the degree by which light onset changes the spontaneous firing rate and to avoid an overestimation of the electrodes recording low spontaneous activity, a response ratio was calculated. The calculation is based on the well-established bias index used to quantify physiological light responses (83, 84) and is analogous to the Michelson contrast used to quantify the contrast in visual images:

Response ratio = (Firing rate _LIGHT ON_ – Firing rate _spontaneous_) / (Firing rate _LIGHT ON_ + Firing rate _spontaneous_).

### Flow-sorting of the cone photoreceptors

A protocol from Palfi et al. (85) was used to dissociate retinal cells. Briefly, PN19-26 control and TSA-treated retinal explant cultures were removed from membranes and incubated in trypsin (Sigma-Aldrich) solution for 20 min at 37°C. Following incubation with trypsin inhibitor (Sigma-Aldrich) cell suspension was washed with HBSS and passed through a 40-µm filter before fluorescence activated sorting (FACS). One biological replicate included at least 2 retinal explants prepared from different animals. Three independent biological replicates from control and TSA-treated retinal explants were used for cone photoreceptor FACS using an ARIAIII cell sorter (BD Biosciences). The sort was performed with a 100 µm nozzle tip, at a sheath pressure of 20.0 psi, and with purity precision. TNXL positive cone photoreceptors were gated as follows: singlets forward scatter (FSC-A vs FSC-H) / singlets side scatter (SSC-A vs SSC-H) / viable cells (FSC-A vs SSC-A) / TNXL+ cells (FSC-A vs TNXL-A) (gating Supplementary Figure 3A, B). The purity of sorted TNXL+ cells was checked by performing post-sort FACS analysis (Supplementary Figure 3C, D).

### Whole transcriptome sequencing and data analysis

2000 to 5000 frozen sorted-cells were lysed in approximatively 10 µl of lysis buffer and cDNA synthesis was performed using the SMART-Seq v4 Ultra Low Input RNA Kit (catalog 634888, Takara Bio). First-strand cDNA synthesis was performed using 20 to 50% of the input and was followed by full-length double-strand cDNA amplification using 17 PCR cycles. The quality of the resulting cDNA was validated using Bioanalyzer and High Sensitivity DNA Kit (Agilent), as well as Qubit dsDNA HS fluorometric quantification (ThermoFisher Scientific). NGS libraries were prepared using 150 pg of cDNA input and the Nextera XT DNA Library Preparation Kit (catalog FC-131-1024, Illumina) with 11 cycles of PCR. Libraries were sequenced as single reads (75 bp read length) on a NextSeq500 (Illumina) with a depth of >20 million reads. Library preparation and sequencing procedures were performed by the same individual and a design aimed to minimize technical batch effects. Data quality of raw RNA-seq reads in FASTQ files was assessed using ReadQC (ngs-bits version 2018_04) to identify potential sequencing cycles with low average quality and base distribution bias. Reads were preprocessed with Skewer (version 0.2.2) and aligned using STAR (version 2.5.4a), allowing spliced read alignment to the mouse reference genome (Ensembl Mus musculus GRCm38). Alignment quality was analyzed using MappingQC (ngs-bits version 2018_04) and visually inspected with Broad Integrative Genome Viewer (version 2.4.0). Based on the genome annotation ITAG (Ensembl 75), normalized read counts for all genes were obtained using subread (version 1.6.0) and edgeR (version 3.28.0). Raw counts data were processed using iDEP, an integrated web application for RNA-seq data analysis (86). To provide access to RNAseq data, we generated a BioJupies notebook (87) link providing an interactive and visual analysis of all the data (https://amp.pharm.mssm.edu/biojupies/notebook/LjknTl51J). Sequencing data are deposited on GEO with the accession number GSE141601.

For differential gene expression (DEG) analysis, gene counts were filtered to only retain genes with a value > 1 cpm (count per million), in at least two samples within at least one group (control or treated), leaving around 14,400 genes for determination of differential expression in each of the pair-wise comparisons between experimental groups. Differentially expressed genes between treated and control groups were identified using two-tailed permutation FDR based Student’s T-test (FDR <0.05 and 250 randomization). Only transcripts coding for protein sequences were retained for pathway analysis. Quantitative gene expression data was submitted and integrated in the PaintOmics 3 data analysis platform (88, 89), in order to identify trends in pathway modulation following the TSA treatment. A stable link is provided to access and visualize the results (http://www.paintomics.org/?jobID=4CIbSr1eA4).

Quantitative RT-PCR. For qRT-PCR, 2000-6000 cones from two pooled PN19-26 *ex vivo rd1^TN-XL^* explants were sorted in 350 µl RLT buffer (Qiagen) to lyse cells. RNA extractions were performed using an RNeasy Micro Kit (Qiagen), followed by cDNA synthesis using the QuantiTect Reverse Transcription Kit (Qiagen). qRT-PCR reactions were performed in technical duplicates of 4 biological samples of TSA-treated and control retinas, on BioRad CFX96 real-time system using QuantiTect SYBR Green PCR Kit (Qiagen) along with gene specific forward (fwd) and reverse (rev) primers (250 nM). The sequences of the primer sets used are listed in Supplemental table 1. The PCR protocol included 40 cycles of: 94 °C (15 s), 57 °C (30 s) and 72 °C (30 s). Relative mRNA expression of each gene of interest was calculated using ΔΔCT method and GAPDH as a housekeeping gene.

#### Statistics

All data were analyzed using Excel (Microsoft) and GraphPad Prism 6. For each comparison between control and treated groups normal distribution was determined with GraphPad software (D’Agostino & Pearson omnibus and Shapiro-Wilk normality tests). Quantification of *in vivo* survival for every time points, for different positions, was calculated using Mann-Whitney nonparametric test. Overall temporal survival curve shown in Figure 2J was obtained by averaging all the analyzed positions from all the animals per stage.

*Ex vivo* survival was quantified on 5-12 different positions, from 3-5 retinal cross-sections obtained from different positions within a retinal explant. Unpaired, two-tailed *t* tests were used to compare controls with treatments.

Statistical differences for LC3-LAMP1 puncta quantification and qRT-PCR were calculated using Mann-Whitney nonparametric test.

RNAseq raw data were processed and normalized using subread (version 1.6.0) and edgeR (version 3.28.0). Statistical data analysis was carried out using the Perseus tool suite for Omics data analysis (90). Two-tailed, unpaired permutation based FDR Students’ T-test on biological replicates mean difference was applied (FDR<0.05 and 250 randomization). Log2 fold changes, mean differences and p-values are reported in the Supplementary table 2.

Statistical difference of light-induced response ratio between control and treated retinas was calculated using a Wilcoxon rank-sum test.

## Author contributions

MS, AC, RGS, AA, EP, WH, CG, PB, DT performed the experiments. All authors analyzed and interpreted the data. DT conceived the research and wrote the manuscript with an input from all authors. First authors’ position is determined by workload.

## Acknowledgements

We are thankful to Klaudija Masarini, Norman Rieger, Patricia Velasco and Coni Imsand for technical assistance. We are grateful to Botond Roska for inspiring scientific discussions. We thank Irena Stingl for graphical design. This work was supported by the ProRetina foundation, the Kerstan Foundation, Deutsche Forschungsgemeinschaft (DFG TR 1238/4-1), Swiss National Science Foundation (31003A_173008), BMBF (FKZ: 01EK1613E), and GC2018-098557-B-I00 from MCIU/AEI/FEDER, UE. RGS is recipient of a Juan de la Cierva contract from MCIU, Spain. We acknowledge support by Open Access Publishing Fund of University of Tübingen.

