## Supplemental Material for "HDAC inhibition ameliorates cone survival in retinitis pigmentosa mice"

### Supplementary information

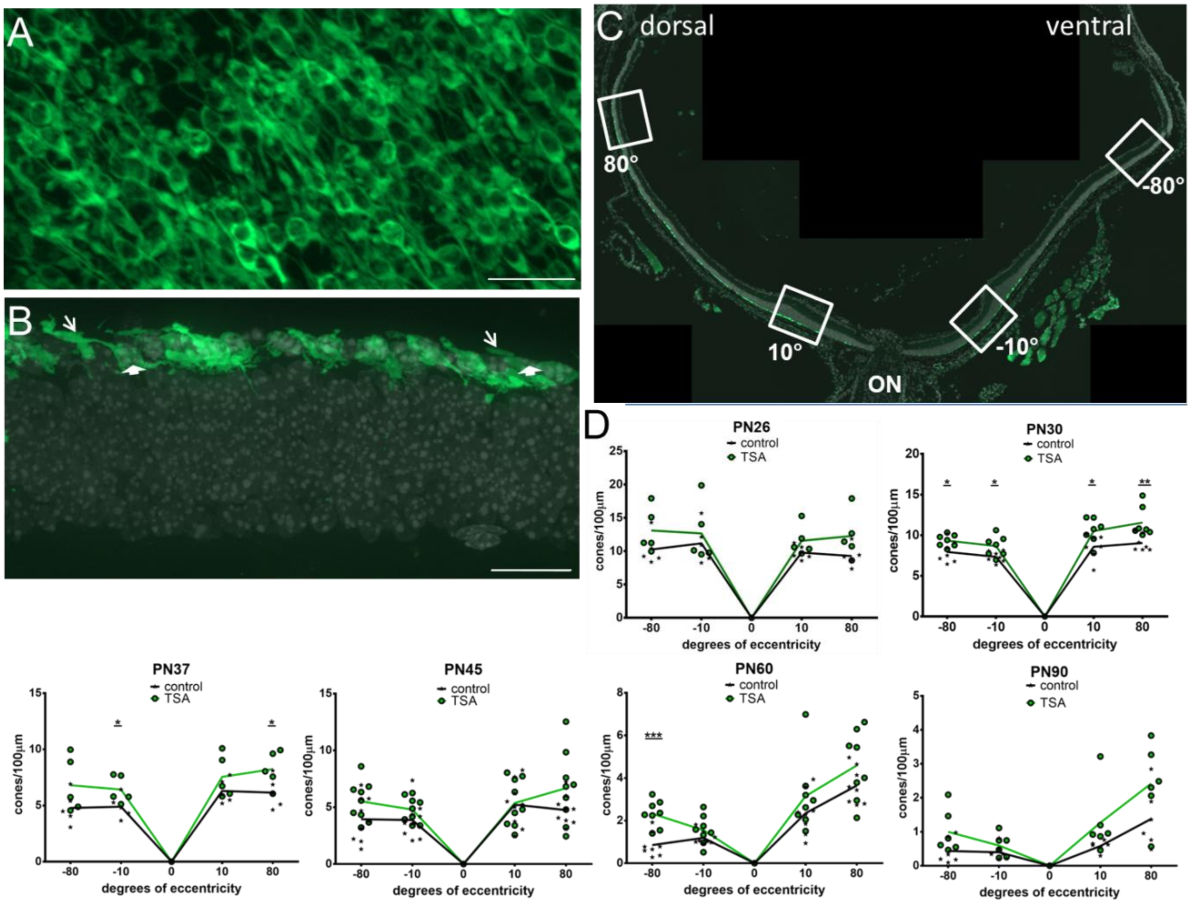


Figure S1. **Quantification of cone photoreceptors.** (A) Representative image of a P30 flat mount *rd1*$^{TN\text{-}XL}$ retina, with TN-XL biosensor labeling cone cell bodies, segments and axons. In the *rd1*$^{TN\text{-}XL}$ retina at PN30 most rods have degenerated, leading to almost horizontal positioning of remaining cone photoreceptors due to the lack of structural support from rods. (B) An example of retinal cross-section from an *rd1*$^{TN\text{-}XL}$ mouse at the same age. DAPI facilitated the distinction of cell bodies (filled arrows) from segments marked with arrows. Only cone cell bodies with clearly visible nuclei were counted. (C) To account for center to periphery gradient of cone loss present in the *rd1*$^{TN\text{-}XL}$ retina, cones were quantified, per 100 µm of ONL length, at two positions in the central retina (ventral and dorsal, corresponding to -10° and 10° of eccentricity from the ON, respectively) and around -80° and 80° covering far periphery. (D) Spider diagrams showing the number of cones at indicated positions in the retina following TSA or sham treatment at P19 and analyzed as indicated. PN26 (n=5 retinas), PN30 (n=7), PN37 (n=5), PN45 (n=9), PN60 (n=8) and PN90 (n=6). Statistical significance for each individual position was assessed using Mann-Whitney nonparametric test, with * $p < 0.05$, ** $p < 0.01$, *** $p < 0.001$. Scale bars: 50 µm. ON, optic nerve

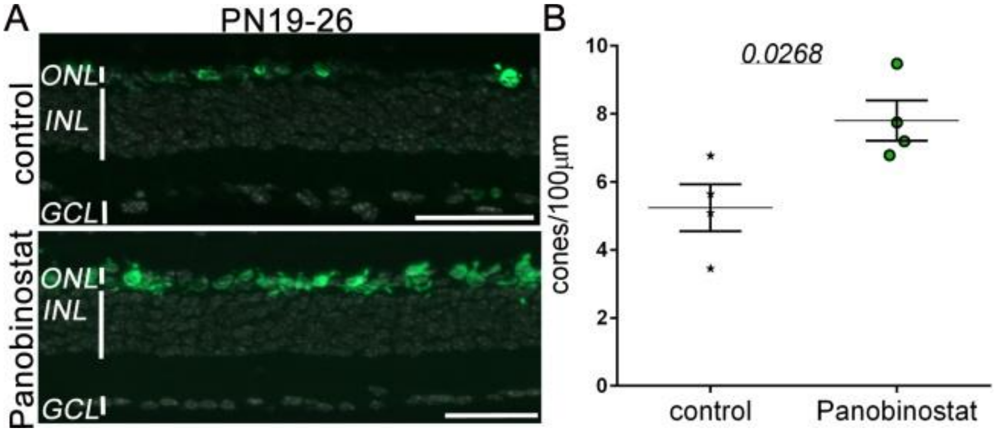


Figure S2. **Panobinostat protects *rd1*$^{TN\text{-}XL}$ cones *ex vivo*.** Retinal explants of *rd1*$^{TN\text{-}XL}$ mice isolated at PN19 were treated with a clinically approved HDAC inhibitor, Panobinostat, for 7 days. (A) Representative images of retinal cross-sections used to count cone number (B). As compared to controls more cones were detected in Panobinostat-treated mice (5.23 ± 0.69 *vs*. 7.8 ± 1.35, respectively). n=4 mice, *p=0.0268,* unpaired, two-tailed t-test. ONL, outer nuclear layer; INL, inner nuclear layer; GCL, ganglion cell layer. Scale bars: 50 µm.

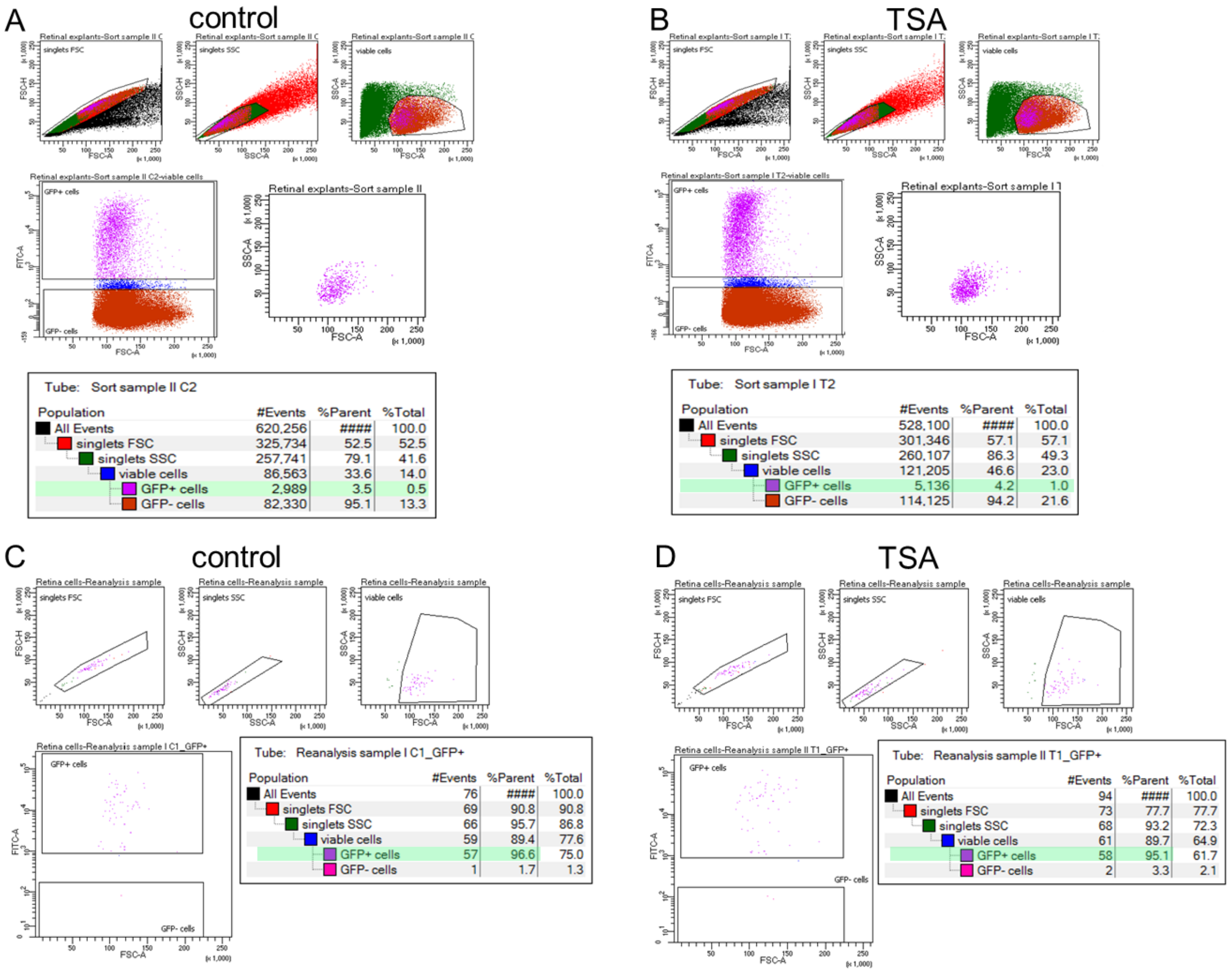


Tube: Sort sample II C2

| Population | #Events | %Parent | %Total |
|---|---|---|---|
| All Events | 620,256 | #### | 100.0 |
| singlets FSC | 325,734 | 52.5 | 52.5 |
| singlets SSC | 257,741 | 79.1 | 41.6 |
| viable cells | 86,563 | 33.6 | 14.0 |
| GFP+ cells | 2,989 | 3.5 | 0.5 |
| GFP- cells | 82,330 | 95.1 | 13.3 |

Tube: Sort sample I T2

| Population | #Events | %Parent | %Total |
|---|---|---|---|
| All Events | 528,100 | #### | 100.0 |
| singlets FSC | 301,346 | 57.1 | 57.1 |
| singlets SSC | 260,107 | 86.3 | 49.3 |
| viable cells | 121,205 | 46.6 | 23.0 |
| GFP+ cells | 5,136 | 4.2 | 1.0 |
| GFP- cells | 114,125 | 94.2 | 21.6 |

Tube: Reanalysis sample I C1_GFP+

| Population | #Events | %Parent | %Total |
|---|---|---|---|
| All Events | 76 | #### | 100.0 |
| singlets FSC | 69 | 90.8 | 90.8 |
| singlets SSC | 66 | 95.7 | 86.8 |
| viable cells | 59 | 89.4 | 77.6 |
| GFP+ cells | 57 | 96.6 | 75.0 |
| GFP- cells | 1 | 1.7 | 1.3 |

Tube: Reanalysis sample II T1_GFP+

| Population | #Events | %Parent | %Total |
|---|---|---|---|
| All Events | 94 | #### | 100.0 |
| singlets FSC | 73 | 77.7 | 77.7 |
| singlets SSC | 68 | 93.2 | 72.3 |
| viable cells | 61 | 89.7 | 64.9 |
| GFP+ cells | 58 | 95.1 | 61.7 |
| GFP- cells | 2 | 3.3 | 2.1 |

Figure S3. **Flow cytometry sorting of TN-XL labeled cone photoreceptors.** (A) FACS plots for a control *rd1*$^{TN\text{-}XL}$ PN19-26 retinal explant. TN-XL positive cone photoreceptors were gated for: singlets forward scatter (FSC-A vs FSC-H) / singlets side scatter (SSC-A vs SSC-H) / viable cells (FSC-A vs SSC-A) / TNXL cells (FSC-A vs TNXL-A, magenta). (B) Plots from a TSA-treated explant. Percentages of fluorescently labeled cones are highlighted in green. (C) The purity of sorted TNXL+ cells was determined by performing post-sort FACS analysis on a resorted control and treated (D) sample.

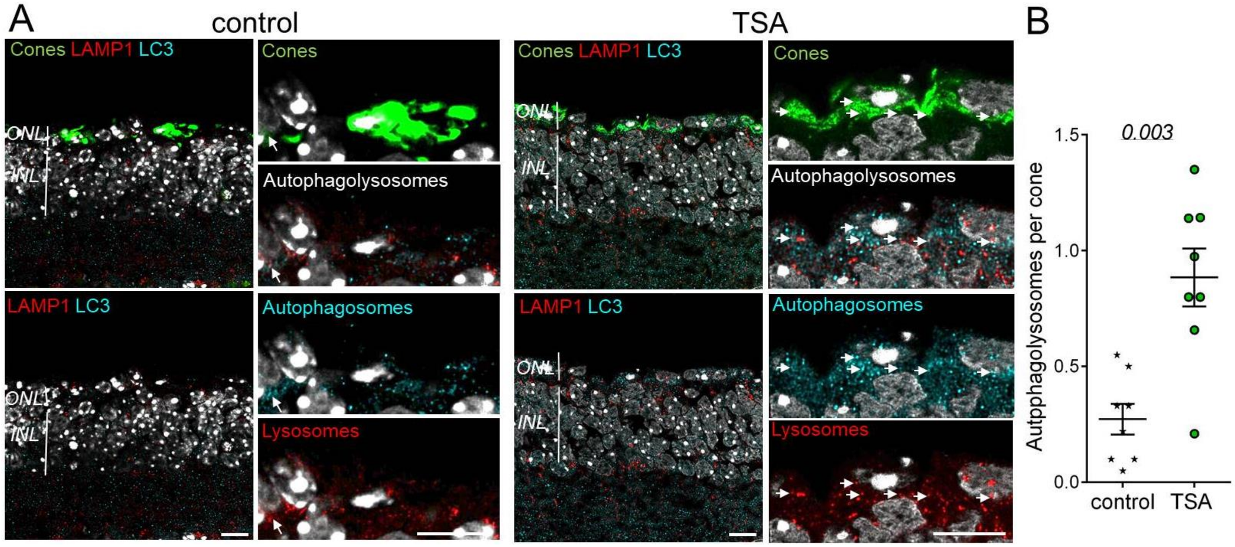


Figure S4. **Autophagy after HDAC inhibition *ex vivo*.** (A) Representative images of a single confocal plane of retinal cross-sections from the *rd1*$^{TN-XL}$ retina explanted at PN19 and treated *ex vivo* for 7 days with control or TSA-medium. Arrows label colocalization between autophagosomes stained with LC3 in cyan and lysosomes labelled with LAMP1 in red. (B) Quantification of the number of colocalized puncta per cone in 4 sections obtained from 2 animals per condition. Data are shown as mean ± SEM. Numerical p-values by Mann-Whitney nonparametric test. Scale bar 10 μm. ONL, outer nuclear layer; INL, inner nuclear layer.

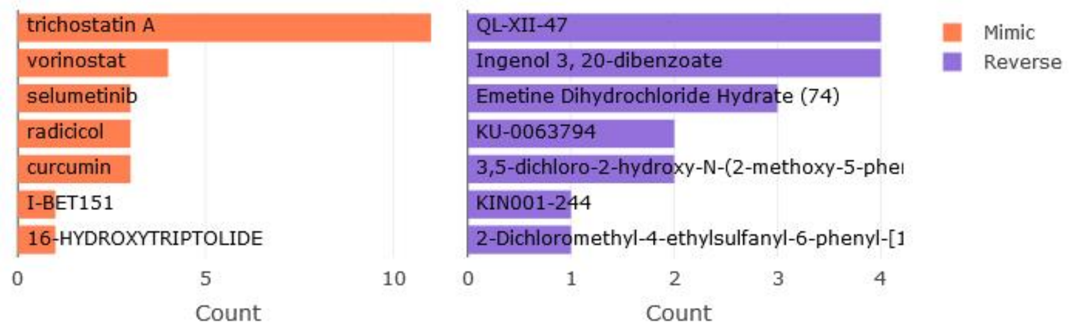


Figure S5. **LINCS Query.** Web-based tool used for identification of small molecules mimicking and reversing the gene expression changes in TSA-protected cones, confirmed the specificity of the observed changes to HDAC inhibitors, Trichostatin A (TSA) and Vorinostat.

Table S1. **Primers used for qRT-PCR analysis.**

| Gene | Forward primer (5’-3’) | Reverse primer (5’-3’) |
|---|---|---|
| *Igf1* | TCCAGTTGCTCTAAGTTTCTCTC | CGTGGGAAGAGGTGAAGATAAG |
| *Bdnf* | TTCGGCCCAACGAAGAAA | TCCTCCAGCAGAAAGAGTAGA |
| *Mtor* | GAGTCCCTCATCAGCATTAACA | ACTCATGCAGCTTCTCATACC |
| *Mlst8* | CCCTGATTCCACGCTTCTT | TGACTCTCCAGGGTTACTACTC |
| *Atg5* | CAAGCCAAGGAGGAGAAGATT | TGCATTTCACGAGAAGAGGAG |
| *Gapdh* | GGAGAAACCTGCCAAGTATGA | TCCTCAGTGTAGCCCAAGA |
